# BMP4 patterns Smad activity and generates stereotyped cell fate organisation in spinal organoids

**DOI:** 10.1101/511899

**Authors:** Nathalie Duval, Célia Vaslin, Tiago Barata, Stéphane Nédélec, Vanessa Ribes

## Abstract

Bone Morphogenetic Proteins (BMP) are secreted regulators of cell fate in several developing tissues. In the embryonic spinal cord, they control the emergence of the neural crest, roof plate and distinct subsets of dorsal interneurons. Although a gradient of BMP activity has been proposed to determine cell type identity in *vivo*, whether this is sufficient for pattern formation in *vitro* is unclear. Here, we demonstrate that exposure to BMP4 initiates distinct spatial dynamics of BMP signalling within the self-emerging epithelia of both mouse and human pluripotent stem cells derived spinal organoids. The pattern of BMP signalling results in the stereotyped spatial arrangement of dorsal neural tube cell types and concentration, timing and duration of BMP4 exposure modulate these patterns. Moreover, differences in the duration of competence time-windows between mouse and human account for the species specific tempo of neural differentiation. Together the study describes efficient methods to generate patterned subsets of dorsal interneurons in spinal organoids and supports the conclusion that graded BMP activity orchestrates the spatial organization of the dorsal neural tube cellular diversity in mouse and human.

## Introduction

The rise of methods to generate pluripotent stem cells (PSC) derived organoids, containing multiple cellular subtypes arrayed in three dimensions (3D) structures, has opened new avenues to decipher the developmental principles underlying the emergence of patterns of cellular differentiation (Huch et al., 2017; Trujillo and Muotri, 2018). These patterns are likely to be established in response to chemical and mechanical cues both externally applied to the organoids through culture conditions and internally generated by the newly created cellular diversity. However, the mechanisms by which cells in organoids interpret these cues and acquire specific cell fates remain largely to be determined.

In order to gain insight into these mechanisms, we focused on cellular subtypes generated in the dorsal part of the embryonic spinal cord (Kalcheim, 2018; Lai et al., 2016). The spinal cord itself originates from cells of the caudal lateral epiblast (or CLE) that transit through a pre-neural (PNP) state before acquiring a full neurogenic progenitor (NP) state (Gouti et al., 2015; Henrique et al., 2015). The CLE and PNP states within this developmental sequence depend on the combined activity of FGF and Wnt signalling, while the transition to the NP state is promoted by retinoic acid (RA) signalling. NP are then directed towards specific neurogenic programmes, depending on their position along the dorso-ventral (D-V) axis of the neural tube (Fig. 1A) (Kalcheim, 2018; Lai et al., 2016). On the dorsal side, NP comprise 6 pools of discrete cell types, named from dorsal to ventral dp1 to dp6 NP, as well as a group of very dorsal cells that will give rise to neural crest cells (ncc) and the roof plate (rp) (Fig. 1A). The dp4 to dp6 NP will give rise to 3 types of early born (dI4 to dI6) and two types of late born (dILA and dILB) associating interneurons (IN), while dp1 to dp3 NP generate dI1 to dI3 relay IN. These neuronal subtypes form functional circuits, in which associating IN transmit the information coming from ncc derived peripheral sensory neurones to relay sensory spinal IN, that in turn convey this information to higher centres in the brain.

**Fig. 1:**
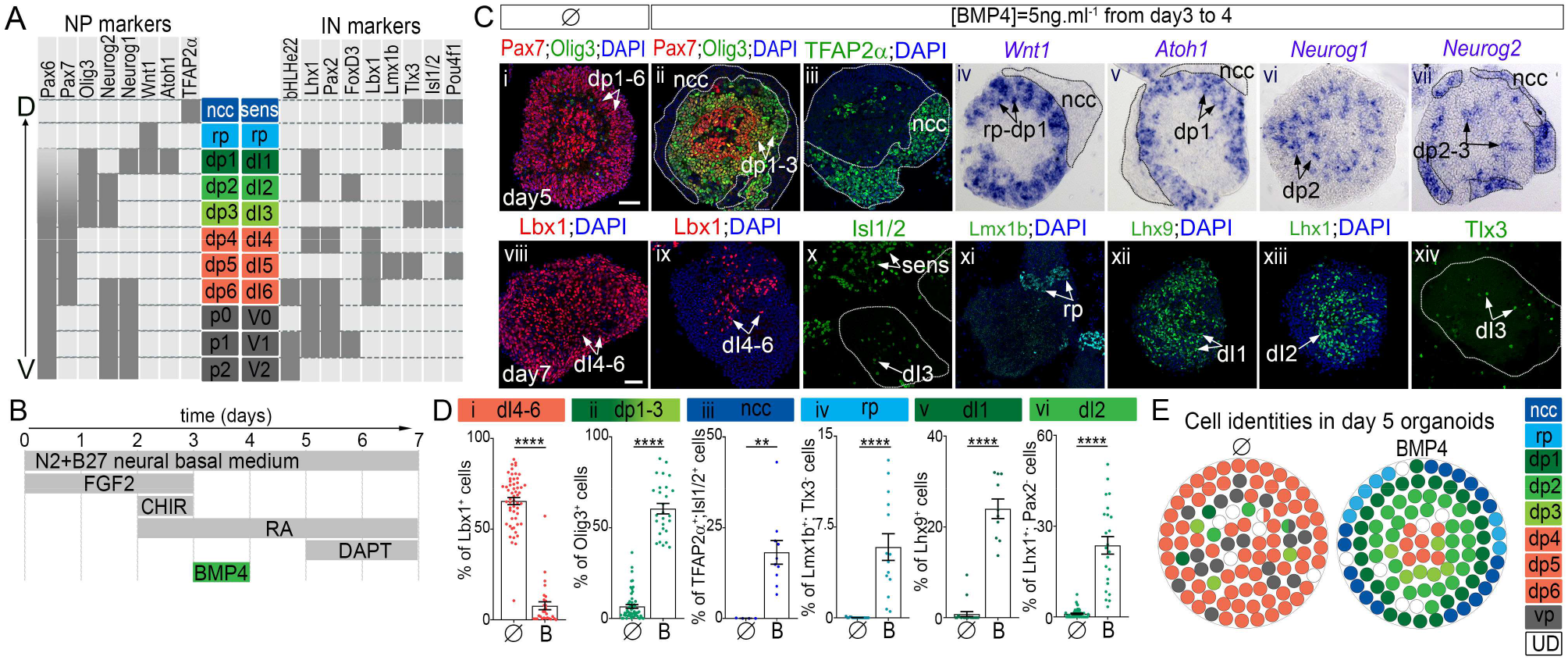
BMP4 patterns mouse ESC derived spinal organoids. **(A)** *In vivo* D-V expression patterns of progenitor (PN) and interneuron (IN) TF markers. **(B)** Schematic representation of differentiation conditions generating dorsal spinal cellular subtypes. **(C)** Immuno-detection **(i-iii, viii-xiv)** and ISH **(iv-vii)** for the indicated dorsal neural cell types. White dash lines in ii-v,vii surround ncc territory, red dash line in ii indicates cells less responsive to BMP4, the white ones in x,xiv delineate EB contour. Scale bars: 60 μm. **(D)** Percentage of specific neuronal cell types harbouring the indicated TF code per image field (circles: individual values, bars: mean±s.e.m.). **(E)** Graphs displaying the distribution and percentage of cell subtypes in day 5 organoids color-coded as in A; white circles mark cells with undetermined fate (UD). sens: peripheral sensory neurons; ncc: neural crest cells.

The commitment of NP towards these discrete dorsal fates is determined by the activity of specific combinations of transcription factors (TF), that display stereotypic temporal and spatial profiles of expression (Fig. 1A)(Kalcheim, 2018; Lai et al., 2016). The profiles of TF marking the ncc, rp, dp1 to dp3 cells are in part generated in response to gradients of diffusing Bone Morphogenetic Proteins (BMPs) secreted first from the non-neural ectoderm, the precursors of ncc and rp cells, then by the rp cells (Kalcheim, 2018; Le Dréau and Martí, 2013; Zagorski et al., 2017). Studies using classical developmental approaches led to the model in which the concentration, timing, and duration of BMP exposure are interpreted by PNP and NP cells to guide them towards a specific fate. Submitting PNP explanted from chick embryos to increasing BMP4 concentrations or exposure durations triggers progressive transcriptional switches and the generation of more and more dorsal cell types (Liem et al., 1995; Sasai et al., 2014; Tozer et al., 2013). The *in vivo* relevance of the levels of BMP signalling in providing positional information has been shown by applying discrete levels of BMP receptor activity to NP in chick embryos (Timmer et al., 2002; Zechner et al., 2003). Similarly, inhibiting BMP signalling using Smad6, an intracellular pathway antagonist, at discrete developmental stages reinforced the idea that duration of BMP signalling discriminates between dorsal cell types (Tozer et al., 2013). Moreover, PNP appears to harbour time regulated competence for dorsal fates specification in response to BMPs. Notably, the ability of the PNP to generate ncc in response to BMP4 is limited in time (Nitzan et al., 2016; Sasai et al., 2014).

However, recent work based on PSC-derived spinal organoids have questioned the morphogenetic potential of BMPs (Andrews et al., 2017; Gupta et al., 2018; Meinhardt et al., 2014; Ogura et al., 2018). Indeed, although exposure to BMPs triggered the dorsalization of PNP or NP, whatever the concentrations of BMPs used, many dorsal neural tube cellular subtypes were generated together in a non-organized fashion in the organoids. This raised the possibility that BMPs act as permissive rather than instructive signals during cell fate acquisition in the dorsal neural tube (Andrews et al., 2017) and that additional signals or mechanisms are responsible for the spatial organisation *in vivo*.

To tackle this issue, we first established robust embryoid body (EB) based protocols to drive mouse or human PSC into PNP that finally acquired a dp4 to dp6 state. We showed that in these organoids, BMP4 created concentric patterns of several dorsal neural tube cell types. These patterns stem from position specific temporal profiles of BMP transcriptional effectors Smad1/5/9 activity within the emerging organoid epithelia. The concentration, duration and timing of exposure to BMP4 modify the content and pattern of cell types produced in these organoids. Furthermore, our data revealed that across evolution the duration of time windows for which PNP and NP are competent to generate specific cell fate in response to BMP4 were modified to match with species specific temporal sequence of neural differentiation. Together, the protocols to generate specific subsets of relay spinal neurons from pluripotent stem cells will serve as useful platforms to dissect the principles underpinning their development and differentiation.

## Material and methods

### Cell lines maintenance and differentiation

Mouse ESC line HM1 (Selfridge et al., 1992) at passage ranging from 15 to 19 were maintained on mitotically inactive primary mouse embryo fibroblasts in EmbryoMax D-MEM supplemented with 10% ESC-qualified fetal bovine serum (Millipore), L-glutamine, nonessential amino acid, nucleosides, 0.1 mM β-mercaptoethanol (Life technologies) and 1000 U.ml^-1^ leukemia inhibitory factor (Millipore). Human iPSC WTSIi008-A cell and WTSIi002-A line (EBISC, European Bank for Pluripotent Stem Cells) were cultured in E8 medium on vitronectin (Life technologies) as previously described (Maury et al., 2015).

To set mouse ESC embryoid bodies differentiation, cells were trypsinized and placed twice onto gelatinised tissue culture plates to remove feeders. The differentiation was initiated by seeding 5 10^4^ cells.ml^-1^ in ultra-low attachment petri dishes (Corning) and in Advanced Dulbecco’s Modified Eagle/ F12 and Neurobasal media (1:1, Life technologies) supplemented with 1x B27 devoid of Vitamin A and 1x N2 (Life technologies), 2 mM L-glutamine (Life technologies), 0.1 mM β-mercaptoethanol, penicillin and streptomycin (Life technologies). The medium was changed every day from the day 2 of differentiation. Human PSC embryoid bodies differentiation was performed as previously described (Maury et al., 2014); except to match mouse differentiation the medium did not contain LDN193189 nor SB431542. 1.75 10^5^ cells ml^-1^ were seeded in ultra-low attachment 6 well plates (Corning). Medium was changed at day 2, 4, 7, 9 and 11 of differentiation. Several chemical drugs were added to the differentiation media for specific time periods to inhibit or activate key developmental important signalling pathways: 10 ng.ml^-1^ bFGF (FGF2, R&D), 3 μM CHIR99021 (Tocris or Axon Medchem), 10 nM Retinoic Acid (Sigma), 1 ng.ml^-1^ to 15 ng.ml^-1^ BMP4 (R&D), 10 μM DAPT (Stemgent).

### Dissociation of mouse ESC derived neuroprogenitors for monolayer culture

EB at day 3 of differentiation were incubated with 0.05% Trypsin-EDTA for 3 minutes. After SVF trypsin inactivation they were dissociated by pipetting. After filtration (40 μm pore size), single cell suspension was plated at 10^5^ cells.well^-1^ in a 24 well plate onto a glass coverslip coated with 20 μg.ml^-1^ poly-ornithine (Sigma) and 5 μg.ml^-1^ laminin (Life technologies). Cells were let to adhere 1h before the addition of fresh differentiation medium containing BMP4.

### Expression Analyses

#### RT-qPCR

Total RNA was extracted from 50 to 250 EB collected at different time points using the NucleoSpin^®^ RNA kit (Macherey-Nagel) following manufacturer’s instructions. cDNA were synthetized using SuperScript IV (Thermo Fischer Scientific), random primers and oligo dT. For real time quantitative PCR (RT-qPCR), SYBR Green I Master (Roche) and the LightCycler 480 II (Roche) were used. PCR primers were designed using Primer3 software (Table S1). Levels of expression per gene for a given time point was measured in biological duplicates or triplicates. Mouse gene expression levels were expressed relatively to *TATA-box Binding Protein (TBP)* mRNA levels and normalized to the expression in either GD11.5 dissected spinal cord, ESCs.

#### Immuno-fluorescence and in situ hybridization

EB fixation, embedding and cryo-sectioning was previously described (Maury et al., 2014), so were the immuno-labelling and *in situ* hybridization (ISH) protocols (Briscoe et al., 2000; Yamada et al., 1993). Details of the reagents are provided in the Table S2. Analyses were carried out using a Leica TCS SP5 confocal microscope or a Zeiss Axioplan 2 and images processed with Photoshop 7.0 software (Adobe Systems, San Jose, CA, USA) or Image J v.1.43g image analysis software (NIH).

#### Quantification

Number of cells immuno-labelled was calculated using Cell Profiler (Broad institute) after nuclei segmentation based on DAPI fluorescence signal and using a signal intensity threshold and was expressed as the percentage of all detected nuclei. The EB area labelled by ISH probes was estimated thanks to Image J v.1.43g image analysis software (NIH) and was graphed as the percentage of the whole EB surface. For each condition, these quantifications were performed on a minimum of 5 images per experiment and on a minimum of 2 independent experiments. Percentage of all cellular subtypes analysed were represented using a dot plot shaped as a circle, position of the cellular subtypes reflects our observations. Levels of PSmads fluorescent signal intensity per cells were evaluated by Cell Profiler and followed by background subtraction evaluated in EB not treated with BMP4. Violin plots were used to show the dispersion of these levels in 5 images per experiments and on a minimum of 3 independent experiments. The fluorescence intensity of PSmads expression along EB outer-inner axis was measured in rectangles of 16 μm wide positioned perpendicular to a EB tangent line with Image J v.1.43g. Background measurements were obtained from EB not treated by BMP4 and these were subtracted from each assayed profile. Statistical analysis and graphs were performed using Prism Graphpad software. Non-parametric t-tests were used to evaluate pair-wise comparisons between conditions. P-values are indicated as * if P ≤ 0.05, ** if P ≤ 0.01, *** if P ≤ 0.001 and **** if P ≤ 0.0001.

## Results and discussion

### Efficient generation of associating IN within spinal organoids

In order to generate dorsal spinal neurons in mouse embryonic stem cells (ESC)-derived organoids, we have adopted 3D embryoid body (EB) based differentiation and adapted culture conditions previously used for the targeted differentiation of PSC into ventral spinal cells (Gouti et al., 2014; Maury et al., 2015; Wichterle et al., 2002). EB were produced in neural basal medium supplemented with FGF2 for the first 3 days of culture and CHIR to activate Wnt signalling between day 2 and 3 (Fig. 1B). However, in contrast to ESC differentiated in a monolayer of cells (Gouti et al., 2014), EB cells treated with these two compounds adopted a mesodermal fate (Fig. 1SA, B, D). Addition of RA on day 2 directed cells towards a neural state (Fig. 1SA, B, D) (Gouti et al., 2015; Henrique et al., 2015). In these conditions and consistent with previous studies, NP harboured a spinal state reminiscent to that found at the cervical and brachial levels of embryos (Fig. S1C, E). Finally, in line with *in vivo* studies (Alvarez-Medina et al., 2008; Lee and Deneen, 2012; Valenta et al., 2011; Zechner et al., 2003), CHIR mediated Wnt signalling activation favoured a dorsal fate acquisition, so that 80% of cells exhibited a dp4 to dp6 molecular identity (Fig. 1Ci, E, S2A, Bv,ix). Accordingly, at day 7 after a 48 hour-long treatment with Notch signalling inhibitor DAPT (Fig. S3A), EB contained mainly neurons reminiscent of the dI4 to dI6 early born associating IN (Fig. 1C viii, Di, S3Bi-iv, C). Thus, the identified combination of FGF2, Wnt agonist, RA, and DAPT represents a fast and efficient means to produce mouse ESC derived organoids containing spinal associating progenitors or neurons (Andrews et al., 2017; Meinhardt et al., 2014).

### Concentric patterns of dorsal neuronal cell types formed in response to BMP4 exposure

We next examined the effects of BMP4, one of the BMP ligands implicated in neural patterning (Le Dréau and Martí, 2013), by exposing day 3 NP for 24h to 5 ng.ml^-1^ BMP4 (Fig. 1B). Consistent with BMP4 being a dorsalising cue, it induced the expression of markers of ncc, rp, as well as relay dp1/dI1 and dp2/dI2 cells (Fig. 1Cii-vii,ix-xiv, Dii-iv, S2Bviii, Ci). Importantly, in day 4 and 5 EB, these cell-type markers displayed stereotyped concentric patterns of expression organised along the outer-inner axis of the organoid, with their relative position matching the one found along the D-V axis of the neural tube (Fig. 1Cii-vii, E). At day 7, the sensory cells produced from the outer ring of ncc detached from the EB to colonize the dish plate (Fig. 1Cx, S3vi, not shown). The rp cells clustered at the periphery of the EB and were surrounded by dI1 IN (Fig. 1Cxi, xii). Inside the EB, dI2 and few dI4-6 associating IN were intermingled (Fig. 1Cix,xiii). Altogether these data demonstrate that in addition to creating cellular diversity, BMP4 directs its spatial rearrangement so that the most dorsal cell types are found at a more peripheral position than ventral ones (Fig. 1E). This organization attenuates overtime due to ncc and post-mitotic IN delamination, which may explain why in previously generated spinal organoids in presence of BMPs and analysed solely for terminally differentiated neurons, the patterning activity of BMPs hasn’t been observed (Andrews et al., 2017; Gupta et al., 2018; Ogura et al., 2018).

### Spatial and temporal dynamics in Smad1/5/9 activity triggered by BMP4 exposure

We next sought to investigate whether BMP4 mediated EB patterning could stem from differential spatial and temporal activation of its intracellular downstream effectors. For this, we monitored the levels of the phosphorylated forms of the TFs Smad1/5/9 (PSmads) from day 3 to 4 in presence or not of BMP4 (Fig. 2A-C).

**Fig. 2:**
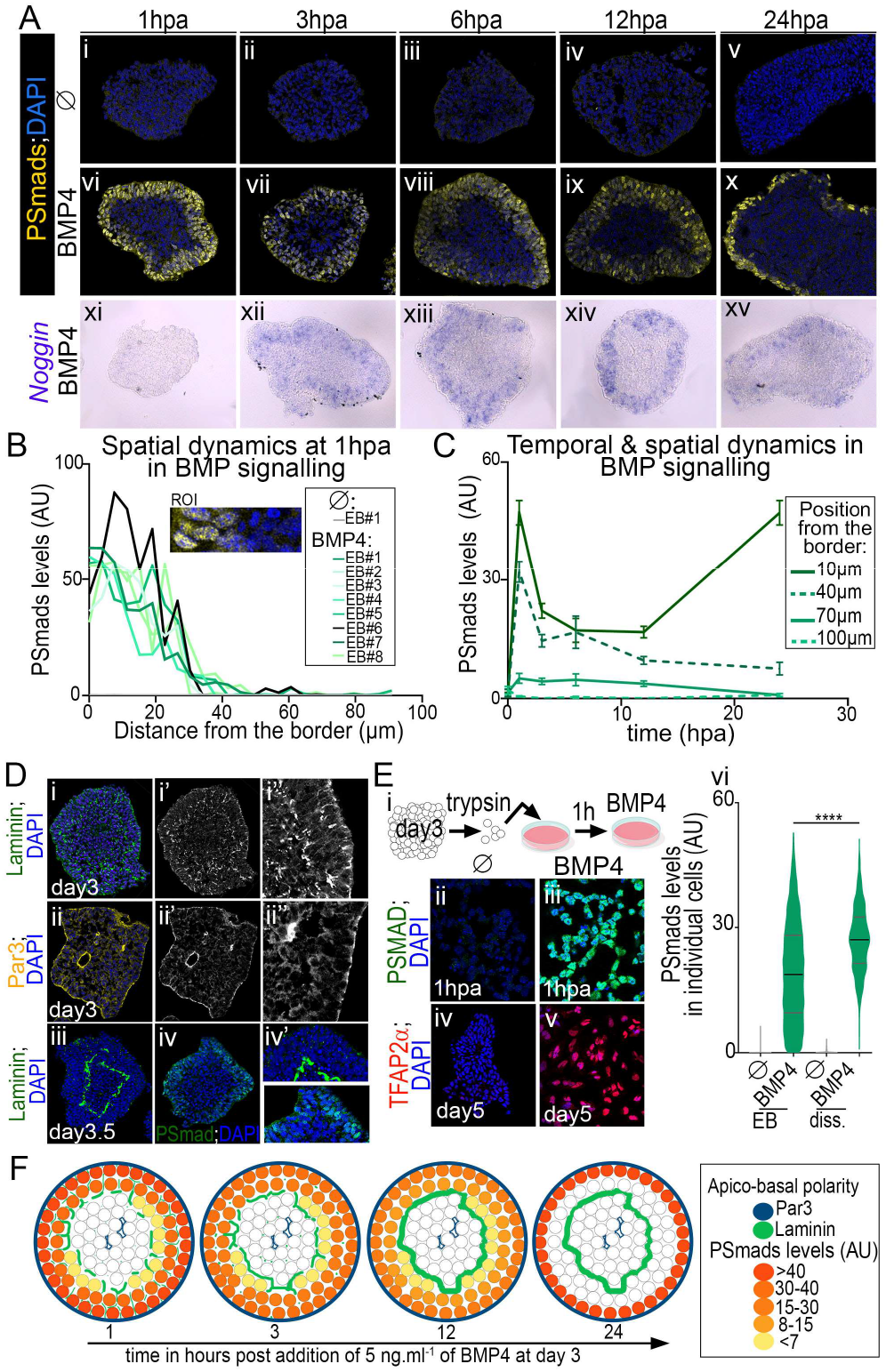
Spatial and temporal dynamics of PSmads in response to BMP4 exposure. **(A)** Immuno-detection of the phosphorylated forms of Smad1/5/9 (PSmads) and DAPI labelling (i-x) and ISH against *Noggin* **(xi-xv)** in EB cultured without or with 5ng.ml^-1^ BMP4 added from day 3 and harvested 1, 3, 6, 12 or 24 hours post BMP4 addition (hpa). **(B)** Inward gradients of PSmads distribution measured in region of interest (ROI) defined as in inset panel and in 8 independent EB 1hpa. **(C)** Temporal and spatial dynamics in PSmads levels in cells positioned at discrete distances from EB borders (mean ± s.e.m.). **(D)** Laminin, Par3, and PSmads immuno-detection and DAPI staining in day 3 and 3.5 EB. In i’, ii’, the laminin and Par3 stainings have been isolated; i”, ii” are blown up of regions in i’, ii’. **(E) (i)** Schematics of experimental procedure to assess the fate of dissociated cells from day 3 EB and exposed to 5ng.ml^-1^ BMP4 1h post-dissociation. **(ii-v)** Immuno-staining of the indicated markers in dissociated cells 1hpa **(ii-iii)** or at day 5 **(iv-v)**. **(iv)** Violin plots showing the dispersion of PSmads levels in EB cells or dissociated cells 1hpa (bars: mean±quartiles). **(F)** Graphs representing the temporal and spatial dynamics in PSmads levels and polarity marker distribution.

At all time-points, PSmads^+^ cells formed a spatially restricted 3 to 5 cells width ring at the periphery of the EB and cells deep inside the organoid were devoid of PSmads signal (Fig. 2A vi-x, B). The limited rate of BMP proteins diffusion could participate to this spatial restriction (Kicheva et al., 2007; Pomreinke et al., 2017; Zinski et al., 2017). Alternatively, the formation of an extracellular matrix separating an outer epithelium from inner epithelia within the organoid may act as a barrier hampering BMP4 diffusion (Hu et al., 2004; Ma et al., 2017; Perrimon et al., 2012; Plouhinec et al., 2013; Ramirez and Rifkin, 2009; Wang et al., 2008) (Fig. 2D, F). The emergence of epithelia was revealed by analysing the distribution profiles of Par3, a member of the apical protein complexes, and of laminin, a component of the basal lamina (Fig. 2D, not shown). PSmads^+^ cells delineated exactly the outer epithelium whose apical side faced the culture medium (Fig. 2D, F). The polarity of cells, which is inverted compared neural organoids grown in matrigel (Meinhardt et al., 2014), could arise from focal adhesions between the CLE and PNP cells on day 2 (not shown, (Bedzhov and Zernicka-Goetz, 2014). Furthermore, treating cells that had been dissociated from day 3 EB with BMP4 triggered a homogeneous activation of Smads and drove all cells towards a ncc state (Fig. 2E). These results further support the idea that the emerging 3D structures within organoids restrict spatially BMP4 response (Fig. 2F).

The analysis revealed also that the levels of PSmads within the outer epithelium varied in space and time. Smads activation displayed a decreasing inward gradient across this epithelium (Fig. 2B). In addition, the amplitude of this gradient decreased progressively overtime (Fig. 2C), except in the outer most layer of ncc cells, where after the initial decline, Smads activity elevated again (Fig. 2Ax,C, F, data not shown). The overall temporal adaptation of BMP signalling was supported by the progressive decrease in the levels of BMP4 target genes (Fig. 2Axv, S2Cii) and is likely to be mediated by negative feedback (Bier and De Robertis, 2015). In agreement, BMP antagonist *Noggin* was transcriptionally induced by BMP4 between 1 and 3 hours post BMP4 addition, when the decline in PSmads levels was observed (Fig. 2Axi-xv). Together, these data indicate that the intracellular response to BMP4 generates stereotyped temporal profiles depending on the position of cells within the organoid epithelium (Fig. 2C, F). This spatial organisation of Smads activity provide an explanation for why the use of BMP4 is not sufficient to drive cells in organoids towards a unique cell type (Andrews et al., 2017; Gupta et al., 2018) but rather creates by itself an organised cellular diversity. Furthermore, the match between this position specific-dynamics and the emergence of distinct cell types argues for a model where “positional information” results from a combination of both the levels and the duration of PSmads (Tozer et al., 2013).

### Three tuneable morphogenetic parameters of BMP4 exposure discriminate between dorsal cell types and enrich organoids with specific relay IN subtypes

To test directly the morphogenetic potential of BMP4 in organoids, we sought to modulate the three main parameters known to influence cell response to morphogens (Sagner and Briscoe, 2017): the ligand concentration, the duration of exposure, as well as the time at which the ligand was added (Fig. 3, S4).

**Fig. 3:**
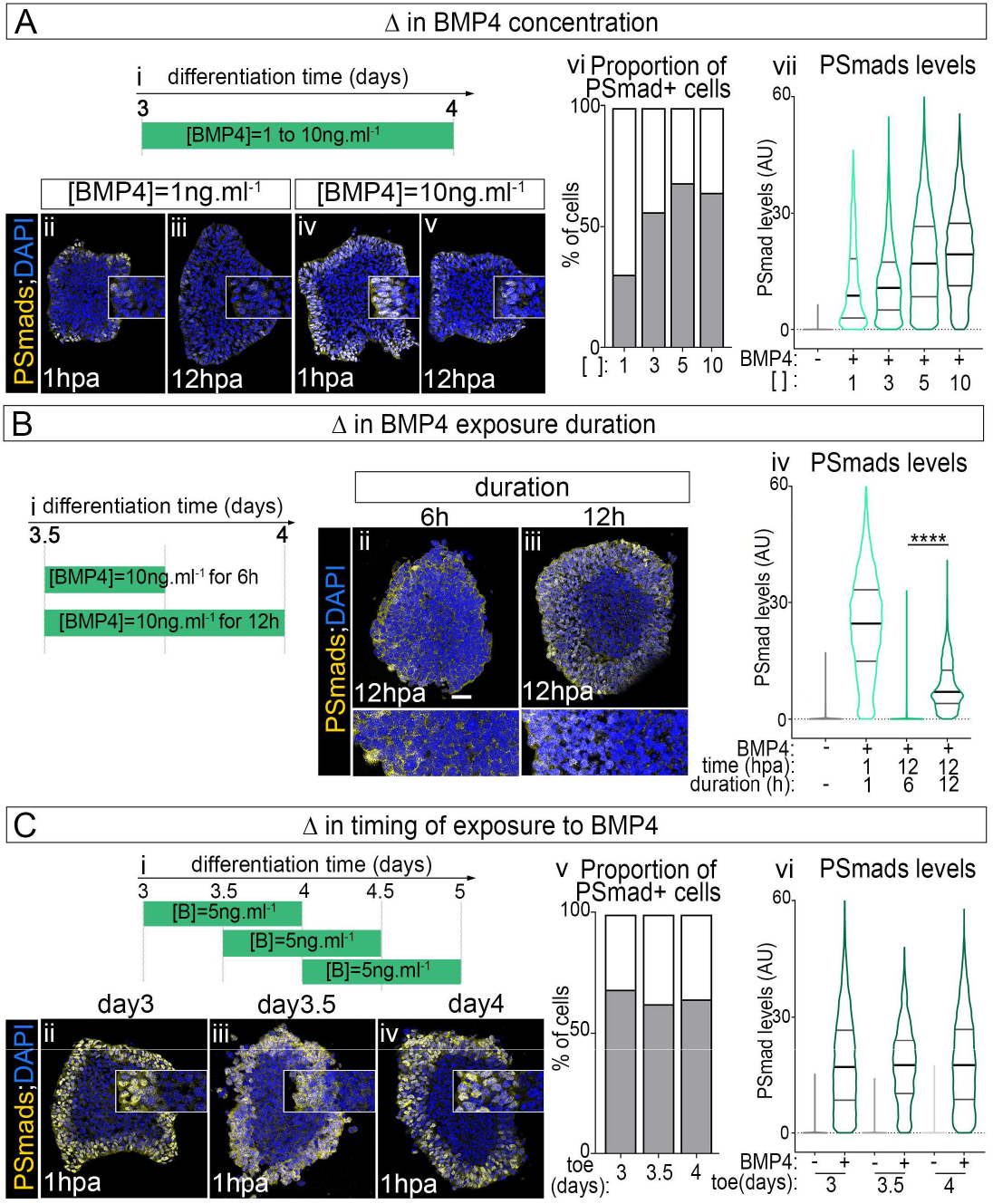
Effects of BMP4 concentration, exposure timing and duration on Smads activity. Characterization of modulating BMP4 concentration **(A)**, BMP4 exposure duration **(B)** and the exposure timing **(C)** on the levels, the spatial distribution and temporal dynamics in PSmads. **(Ai, Bi, Ci)** Schematics showing the time periods of exposure to BMP4 and BMP4 concentrations used in the aside experiments. **(Aii-v, Bii,iii, Cii-iv)** Immuno-detection of PSmads and DAPI labelling in EB at the indicated time (hpa; hour post BMP4 addition) in the indicated conditions of exposure; insets in A and C, lower panels in B are blown ups on PSmads^+^ region. Scale bars: 60 μm. **(Avi, Cv)** Proportion of PSmads^+^ cells per image field. **(Avii, Biv, Cvi)** Violin plots showing the distribution of cells as a function of PSmads levels in organoids in the indicated BMP4 exposure condition and time of analysis.

Increasing BMP4 concentration enlarged the number of PSmads^+^ cells (Fig. 3Avi), the mean levels in PSmads (Fig. 3Avii), as well as the period of time for which the Smads were active (see insets in Fig. 3Aii-v). Similarly, the duration of BMP4 exposure altered the temporal dynamics of BMP4 signalling. The levels of nuclear PSmads ceased as soon as 6 hours after BMP4 removal (Fig. 3B), arguing against a long-term memory of intracellular signalling by cells exposed to BMP (Tozer et al., 2013). Importantly, the modulation of PSmads dynamics by increasing BMP4 concentration or duration extended the total amount of BMP dependent cell-types generated and promoted more dorsal cell types at the expense of ventral ones (Fig. 4A, C, D, S4A, B).

**Fig. 4:**
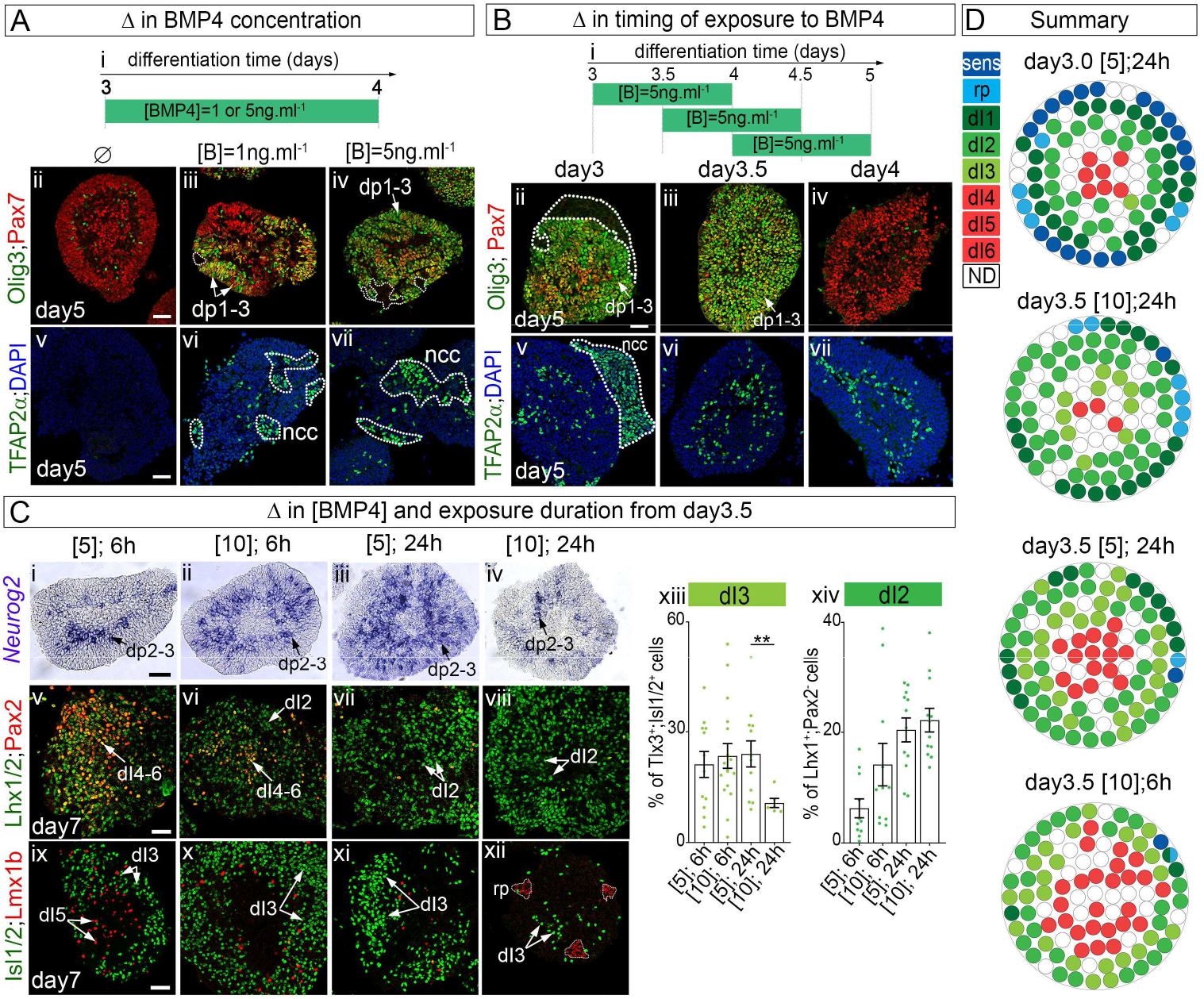
Tuneable morphogenetic parameters of BMP4 exposure enriching organoids with specific relay IN subtypes. Phenotypic characterisation of EB upon the modulation of BMP4 concentration **(A)**, of BMP4 exposure timing **(B)** and of both BMP4 concentration and exposure duration adding BMP4 at day 3.5 **(C)**. **(Ai, Bi)** Schematics indicating when cells are exposed to BMP4 and at which concentration. **(Aii-vii, Bii-vii, Cv-xii)** Immuno-staining for the indicated markers in EB grown in the indicated conditions at either day 5 or day 7. **(Ci-iv)** ISH for the *Neurog2* in day 5 EB cultured in the indicated conditions. Scale bars: 60 μm. **(Cxiii, xiv)** Percentage of dI2 and dI3 IN harbouring the indicated TF code per image field (circles: individual values, bars: mean ± s.e.m.). **(D)** Graphs displaying the distribution and percentage of cell-subtypes in day 7 organoids cultured in the indicated conditions; white circles mark cells with undetermined fate (UD).

Changing the time at which EBs were exposed to BMP4 also had marked effects on the cell fate adopted (Fig. 4B, S4C and compare Fig. 1C, D to Fig. 4D). Cells displayed a 12 hours competence time window for generating specific BMP4 dependent cell types, the most dorsal cell types requiring an earliest time of exposure than more ventral ones (Fig. 4Bii-vii). This is in agreement with the limitation in time for chick spinal NP to become ncc in response to BMP4 (Sasai et al., 2014). Importantly, the switches in cell competence were not due to alteration in the ability of cells to transduce BMP4 information intracellularly. The amplitude (Fig. 3Cvi), gradient (insets in Fig. 3Cii-iv, 3Cv) and temporal adaptation (not shown) of PSmads profiles remained invariant upon shifts in the timing of BMP4 exposure. Instead, the competence could stem from changes in the molecular state of NP during the course of their differentiation (Fig. S2Bi-vi) (Sasai et al., 2014).

Finally, in agreement with concentration, timing and duration of exposure to BMP4 as tuneable morphogenetic exposure parameters (Sasai et al., 2014; Tozer et al., 2013), we identified conditions to obtain organoids containing a dominant relay IN subpopulation at their periphery (Fig. 4C,D), including dI2 and dI3. Previous protocols had reported poor yields of these cell types (Andrews et al., 2017).

### Evolutionary alterations in the length of NP competence time windows to accommodate with discrete tempo of neuronal differentiation

Dynamics of neural differentiation and tissue sizes are highly variable between species (Ebisuya and Briscoe, 2018), raising the question of whether morphogen interpretation is modulated to scale with these dynamics. To address this question, we have adapted our protocol to generate ventral spinal neurons from human iPSC, by removing the Shh agonist SAG (Maury et al., 2014) (Fig. 5Ai). As for mouse ESC, this protocol was sufficient to generate spinal organoids containing mostly dp4-dp6 NP at day 9 of differentiation and associating neurons at day 14 (Fig. 5Aii,iii). Importantly, while both human and mouse dorsal spinal organoids contained similar number of cells (not shown), the time frame of their differentiation is increased by around 2.5 fold in human compared to mouse. Hence, as for motor neuron generation, the tempo of differentiation is a major difference in dorsal differentiation between vertebrate species (Maury et al., 2015; Wichterle et al., 2002).

**Fig. 5:**
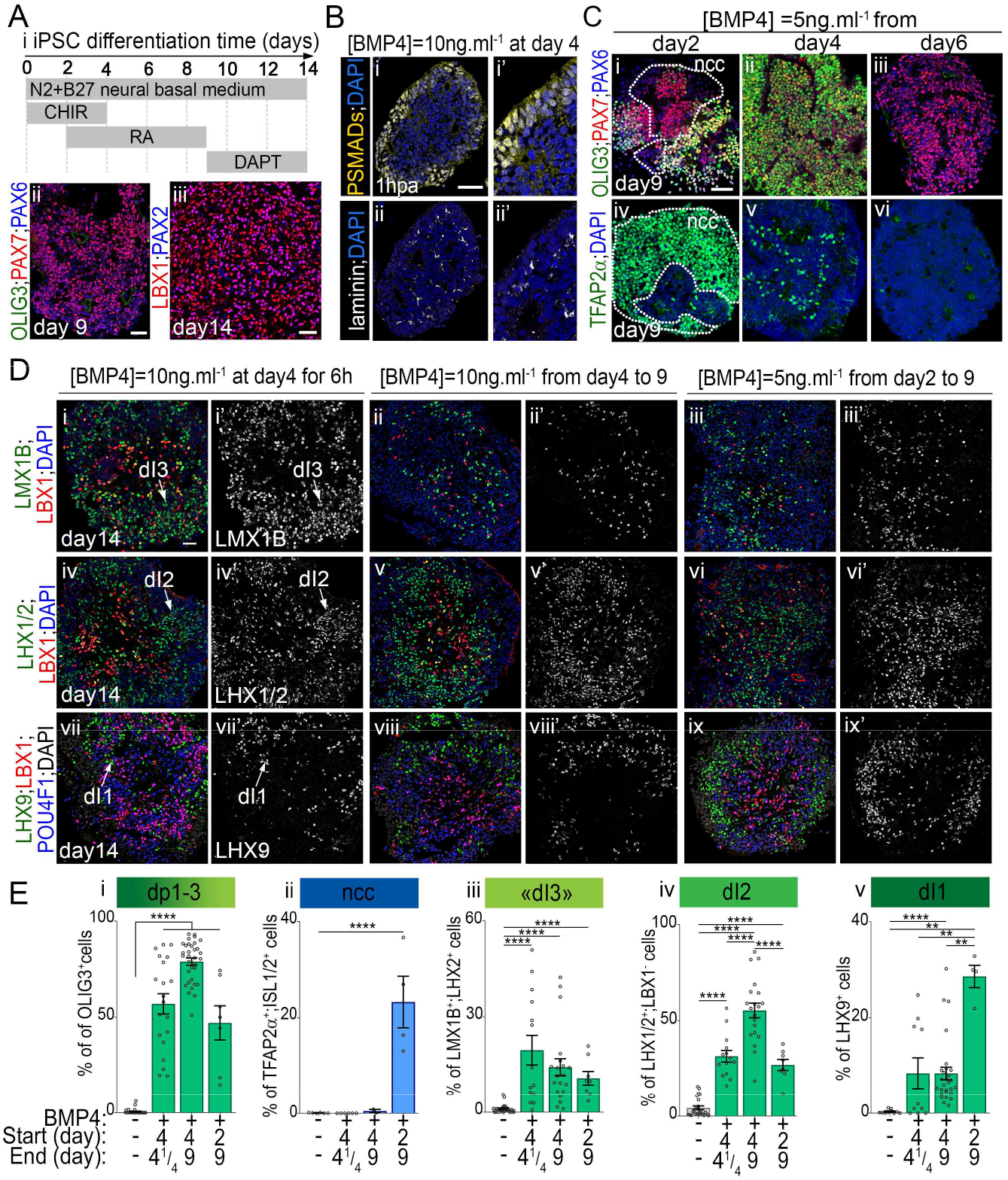
Interpretation of BMP4 exposure by human iPSC derived spinal organoids. **(A)** Generation of associating progenitors and IN from human iPSC. **(Ai)** Schematics indicating the period of drug treatments to convert hiPSC into dorsal spinal cells in organoids. **(Aii-iii)** Immuno-staining for the indicated NP (ii) and IN (iii) markers in day 9 and 14 organoids, respectively. **(B)** Immuno-detection of PSMADs or LAMININ and DAPI labelling in day 4 organoid treated for 1 hour with BMP4. **(C)** Immunostaining for the indicated NP markers in day 9 EB grown in presence of BMP4 added at the indicated time points (day) and maintained up to day 9. **(D)** Immuno-staining for the indicated post-mitotic markers in day 14 EB grown in the indicated conditions. **(E)** Percentage of IN subtypes harbouring the indicated identities (circles: individual values, bars: mean±s.e.m.). Scale bars: 60 μm.

The analysis of the effects of BMP4 on iPSC derived organoids revealed that the response of human cells to BMP4 is similar to that of mouse cells. As with mouse EBs, Smads activation was restricted to the outer layer of cells, which was separated from more inner cells by a basal lamina (Fig. 5B). PSmads levels were the highest in the most peripheral cells and tended to decreased within the width of the epithelium (Fig. 5Bi,i’). Accordingly, several BMP cell types were produced in concentric manner in presence of BMP4 with the most dorsal subtypes been located at a more peripheral position than the ventral ones (Fig. 5D). Finally, as with mouse cells, both BMP4 concentration and the exposure duration influenced the proportion of BMP-dependent cell types produced (Fig. 5D, E). Human NP also displayed competence time windows for the generation of ncc or dp1-dp3 cells (Fig. 5C). Yet, these lasted substantially longer than in mouse (compare Fig. 5C to 4B) and they paralleled the species specific time frame of cell fate acquisition. This brings support to the idea that competence is a consequence of sequential transitions in the transcriptional states of differentiating NP which occur at a slower pace in human (Sasai et al., 2014).

Together our data identified crucial parameters underpinning cell fate switches triggered in organoids treated with BMP4. Firstly, the limited range of BMP4 activity in the emerging 3D structures of the organoids results in the generation of stereotyped profiles of Smads activity in a concentric manner. Changes in the concentration and exposure duration of BMP4 modulate these dynamics in time and space, within the limits imposed by the 3D structures. From these profiles, dorsal progenitor subtypes are spatially organised. This stereotypic arrangement attenuates upon cell delamination and leads to the formation of organoids with intermingled relay IN subtypes. Importantly, these results call for the elaboration of new diffusible compounds capable of activating this pathway in order to be able to reach homogeneity upon targeted differentiation. Secondly, the time of differentiation, i.e the epigenetic state of the receiving cells, appeared as the most constraining parameter to the response of cells to BMP4, with species specificities. The temporal switches in cell competence to respond to BMP4 extend the morphogenetic outputs of this ligand. It also provides robustness to its interpretation, as it no longer relies on the sole precision in diffusion gradient. This certainly explain why dynamics in competence of cells to respond to morphogens is recurrently used during pattern formation (Sagner and Briscoe, 2017). As a consequence, it will require precise temporal control of morphogen signalling by drug treatments during targeted differentiation carried out in organoids.

## Contributions

Conceptualization: V.R., S.N., N.D.; Experiments: N.D., C.V., V.R., T. B.; Writing: V.R., S.N., N.D.; Supervision: V.R., S.N.; Funding acquisition: V.R., S.N.

## Acknowledgement

We deeply thank C. Birchmeier, T. Müller, J.B. Brunet and A. Pierani for antibodies and the ImagoSeine core facility of Institut Jacques Monod, a member of France-BioImaging (ANR-10-INBS-04) and certified IBiSA. We are grateful to Line Manceau, Youcef Frarma, Benoit Sorre and James Briscoe for critical comments on the manuscript.

## SUPPLEMENTARY INFORMATION

### Supplementary material and methods

***Primer sequences usedfor qRT-PCR on mouse EB***

Fw-*Atoh1*: GCTGGTAAGGAGAAGCGGCTGTG

Rev-*Atoh1*: TGTACCCCATTCACCTGTTTGC

Fw-*Fgf5:* TGTACTGCAGAGTGGGCATC (from (Turner et al., 2017))

Rev-*Fgf5:* ACAATCCCCTGAGACACAGC (from (Turner et al., 2017))

Fw-*Hoxb1:* GGTCAAGAGAAACCCACCTAAG

Rev-*Hoxbl:* CACGGCTCAGGTATTTGTTG

Fw-*Hoxb4*: CACGGTAAACCCCAATTACG

Rev-*Hoxb4*: GAAACTCCTTCTCCAACTCCAG

Fw-*Hoxc6*: ACACAGACCTCAATCGCTCAG

Rev-*Hoxc6*: C GAGTT AGGT AGC GGTTGAAG

Fw-*Id2*: CTCCAAGCTCAAGGAACTGG

Rev-*Id2*: TGCTATCATTCGACATAAGCTCAG

Fw-*Klf4:* GGAAGGGAGAAGACACTGCG

Rev-*Kf4*: ATGTGAGAGAGTTCCTCACGCC

Fw-*Neurog1:* GACACTGAGTCCTGGGGTTC

Rev-*Neurog1*: GTCGTGTGGAGCAGGTCTTT

Fw-*Neurog2:* GTGCAGCGCATCAAGAAGAC

Rev-*Neurog2*: TGAGCGCCCAGATGTAATTG

Fw-*Nkx1.2:* ACTGCCTTCACTTACGAGCA

Rev-*Nkx1.2:* AAATTTTGACCTGCGTCT

Fw-*Pou5f1:* AGTTGGCGTGGAGACTTTGC

Rev-*Pou5f1:* CAGGGCTTTCATGTCCTGG

Fw-*TBP*: GAAGAACAAT C C AGAC T AGC AGCA

Rev-*TBP*: CCTTATAGGGAACTTCACATCACAG

**Supplementary Table 1:** List of antibodies

| Antigen | Species | Dilution | Provider (reference) |
| --- | --- | --- | --- |
| **bHLHE22** | Rabbit | 1/2500 | Abcam (ab204791) |
| **Brachury** | Goat | 1/300 | R&D systems (AF2085) |
| **Pou4f1** | Mouse | 1/400 | Chemicon (A5945) |
| **Cdx2** | Rabbit | 1/1000 | Abcam (76541) |
| **Dbx1** | Rabbit | 1/100 | Pierani et al., 1999 |
| **FoxD3** | Guinea pig | 1/5000 | From T. Müller and K. Birchmeier (Muller et al., 2005) |
| **Gsx2** | Rabbit | 1/3000 | Millipore (Gsh2, ABN162) |
| **Hoxa5** | Rabbit | 1/1000 | Sigma (HPA029319) |
| **HuC/D** | Mouse | 1/1000 | Thermo Fisher Scientific (16A11) |
| **Islet1/2** | Mouse | 1/50 | DHSB clone 39.4 |
| **Laminin** | Rabbit | 1/500 | Sigma (L9393) |
| **Lbx1** | Guinea pig | 1/10000 | From T. Muller and K. Birchmeier |
| **Lhx1/5** | Mouse | 1/20 | DSHB (4F2) |
| **Lhx9** | Goat | 1/200 | Santa Cruz (sc19350 E-14) |
| **Lmx1b** | Rabbit | 1/10000 | From T. Muller and K. Birchmeier |
| **Lmx1b** | Guinea pig | 1/10000 | From T. Muller and K. Birchmeier |
| **Nkx6.1** | Mouse | 1/50 | DSHB (F55A10) |
| **Olig3** | Guinea pig | 1/10000 | From T. Muller and K. Birchmeier |
| **Par3** | Rabbit | 1/1000 | Millipore |
| **Pax2** | Rabbit | 1/500 | Thermo Fisher Scientific (716000) |
| **Pax6** | Rabbit | 1/250 | Biologend (PRB-278P) |
| **Pax6** | Mouse | 1/50 | DHSB (AB 528427) |
| **Pax7** | Mouse | 1/50 | DSHB (PAX7) |
| **PSmad1/5/9** | Rabbit | 1/250 | Cell Signalling Technology (13820S) |
| **Sox1** | Goat | 1/200 | R&D systems (AF3369) |
| **Sox2** | Rabbit | 1/200 | Thermo Fisher Scientific (48-1400) |
| **Tlx3** | Rabbit | 1/10000 | From K. Birchmeier |
| *Secondary antibodies:* donkey against mouse, goat, guinea pig or rabbit IgG coupled to Alexa Fluorophores A488, A546, A568 or A647 (Thermo Fisher Scientific) were diluted 1:300 and used with DAPI (500ng/ml, Sigma, 28718-90-3). |  |  |  |

**Fig. S1:**
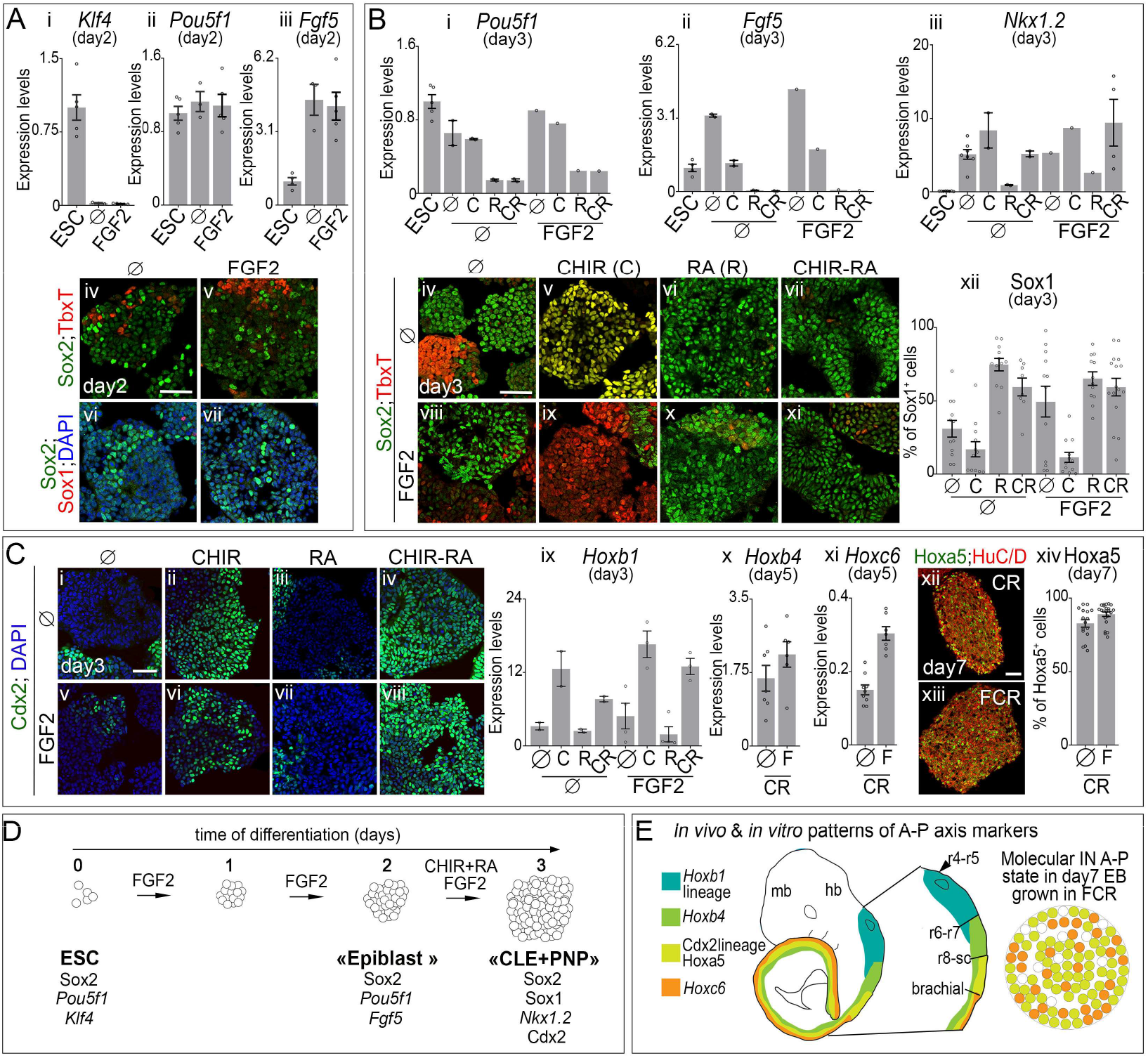
FGF2, CHIR and RA impact on neural and caudal states of mouse ESC derived EB. **(A) (Ai-iii)** Levels of pluripotency markers *Klf4*, *Pou5f1 (Oct4)*, and *Fgf5* expression in ESC or in EB on their 2^nd^ day of differentiation grown without any supplementations (ø) or with FGF2, quantified by RT-qPCR relative to *TBP* and normalized to their levels in ESC (circles: individual values, bars: mean±s.e.m.). **(Aiv-vii)** DAPI labelling and immuno-detection of TbxT, Sox2, and Sox1 in EB after two days in the indicated media. Scale bar:60μM. *By the 2^nd^ day of differentiation, EB cells lost their Pou5f1^+^;Klf4^+^ pluripotency state and acquired epiblast molecular traits (Pou5f1^+^;Fgf5^+^;Sox2^+^), and barely expressed specific lineage markers, such as Sox1 (neural) and TbxT (mesodermal).* **(B) (Bi-iii)** Expression levels of *Pou5f1, Fgf5* and *Nkx1.2* in ESC or in EB on their 3^rd^ day of differentiation quantified by RT-qPCR relative to *TBP* and normalized to their levels in ESC in i and ii and in iii to levels in the tail bud of mouse embryos at gestational day (G.D.) 8.5 (circles: individual values, bars: mean ± s.e.m.). **(Biv-xi)** DAPI labelling and immuno-detection of TbxT and Sox2 in EB after three days in the indicated media. Scale bar: 60μM. **(Bxii)** Quantification of Sox1^+^ cells grown in the indicated medium at day 3 (circles: individual values, bars: mean±s.e.m.). *RA promoted the transition from an epiblast state to a neural state (Sox1^+^; Sox2+) and inhibited the synergic induction of the mesodermal marker TbxT by FGF2 and CHIR. Cells which are not differentiated may harbour a Nkx1.2^+^ CLE state.* **(C) (Ci-viii)** DAPI staining and immuno-detection of Cdx2 after 3 days of differentiation in the indicated media. Scale bar: 60μM. **(Cix,x,xiv)** Levels of *Hoxb1, Hoxb4* and *Hoxc6* quantified by RT-qPCR, relative to *TBP* expression and normalized to their levels in the spinal cord of G.D.11.5 mouse embryos in the indicated media and at the indicated stage of differentiation (circles: individual values, bars: mean±s.e.m.). **(Cxi-xiii)** Immunostaining for Hoxa5 and HuC/D in the indicated conditions at day7 (xi-xii) and quantification of the percentage of Hoxa5+ cells per image field (circles: individual values, bars: mean±s.e.m.; scale bar: 60μM). C, R, F stands for CHIR, RA and FGF2, respectively. *CHIR triggered the caudalization of cells marked by the combined induction of Cdx2, Hoxb1, Hoxb4, Hoxa5 and Hoxc6, FGF2 increased Hoxc6 expression. The molecular identities displayed by cells during their course of differentiation [Cdx2+ at day3, Hoxb4+;Hoxc6+ at day5 and Hoxa5+ at day7] in presence of FGF2, CHIR and RA indicate that they are driven towards a cervical-brachial state (Britz et al., 2015; Dasen et al., 2005).* **(D)** Schematics showing the state transitions of EB cells grown in presence of FGF2 between day0 and 3, CHIR and RA between day 2 and 3. **(E)** Schematics showing the profiles of expression of *Hoxb4*, *Hoxa5* and of *Hoxc6*, as well as the position of cells that had have expressed *Hoxb1* and Cdx2 in GD9.5 mouse embryos. Representation of day7 EB grown in presence of FGF2 between day 0 and 3, CHIR and RA between day 2 and 3 inferred from data presented in D. *Cells in EB grown in these conditions harboured a antero-posterior (A-P) state reminiscent of that found at the cervical-brachial levels of embryos.*

**Fig. S2:**
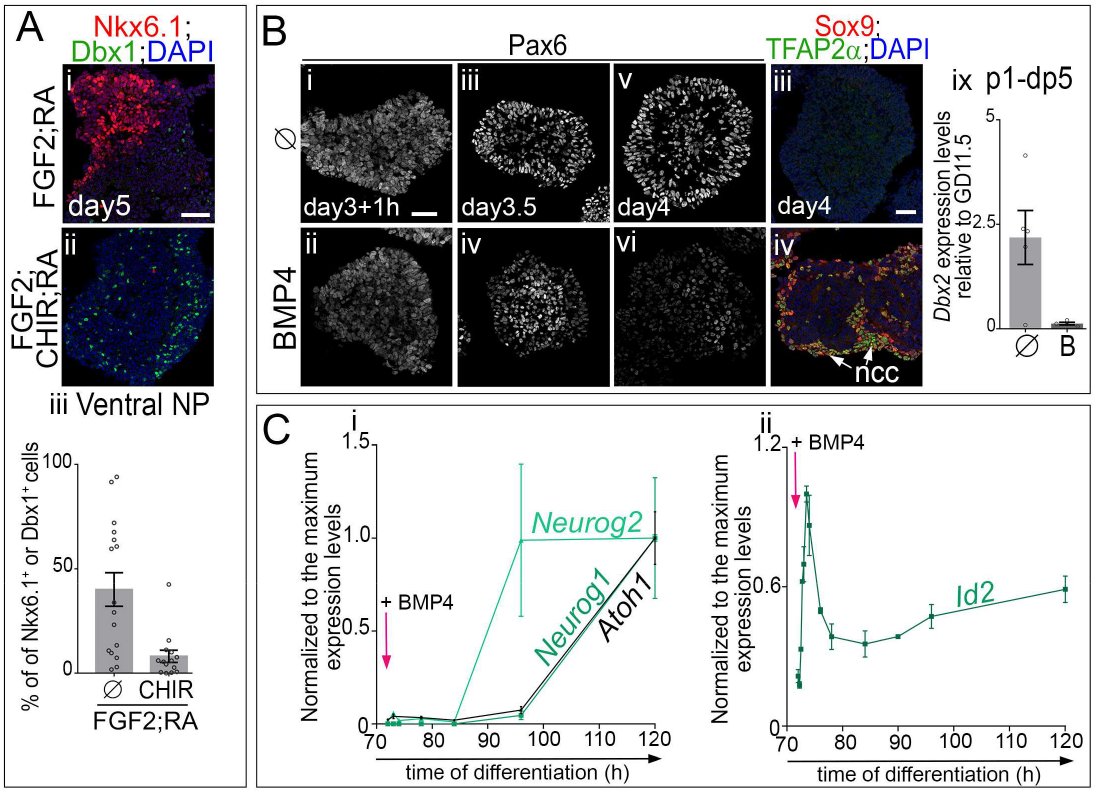
Temporal and spatial dynamics in NP markers expression within dorsal spinal organoids. **(A)** Immunostaining for Nkx6.1 and Dbx1 and DAPI labelling on day 5 EB cultured with FGF2 and RA with (ii) or without (i) CHIR (scale bar: 60μM) and quantification of the proportion of cells expressing either Nkx6.1 or Dbx1 per image field (circles: individual values, bars: mean±s.e.m.). *CHIR mediated Wnt signalling activation dorsalised the states of NP in mouse derived EB, as shown by Nkx6.1 downregulation when CHIR is introduced in the culture medium.* **(B)** Immunostaining for Pax6 or TFAP2ß and Sox9 and DAPI staining on EB at the indicated stages (i-viii; scale bars: 60μM) and expression levels of *Dbx2*, a p1 to dp5 progenitor marker, assessed by RT-qPCR and normalized to its expression levels in G.D.11.5 mouse spinal cord (ix) in day5 EB. In these experiments EB were grown with FGF2, CHIR and RA without or with 5ng.ml^-1^ of BMP4 between day 3 and 4 of culture. *Without BMP4 Pax6 expression increased between day3 and day3.5 suggesting that NP have acquired a new transcriptional state; this could underpin the lost in the ability of cells to acquire a ncc state upon BMP4 exposure from day3.5 (i-iii, see Fig. 4B). In presence of BMP4, Pax6 levels decreased at the periphery of the EB as soon as 12hpa to BMP4, which is in agreement with cells acquiring a more dorsal fate (iv-vi, see Fig. 1A). In presence of BMP4 on day3 of differentiation cells at the periphery of the EB adopted a TFAP2ß; Sox9^+^ ncc fate on day4. The intermediate dorsal spinal cord marker Dbx2 was present in day5 EB grown in absence of BMP4 which is in agreement with most EB cells have adopted a dp4-dp6 fate. Its expression decreased upon BMP4 treatment.* **(C)** Expression levels for *Neurog2* (pale green line in i), *Neurog1* (dark green line in i), *Atoh1* (black line in i) and *Id2* (ii) assessed by RT-qPCR from day 3 to day 5 in EB cultured as in Fig. 1A with BMP4. *In presence of BMP4, cells first induced the dp2/dp3 marker Neurog2 before to induce the dp1 and dp2 markers Atoh1 and Neurog1, this temporal sequence is similar to that found *in vivo* in cells that will acquire a dp1 state (Tozer et al., 2013). The temporal profile of BMP signalling target Id2 confirmed that intracellular BMP signalling progressively decreased as soon as 1h after addition of BMP4 (Samanta and Kessler, 2004; Hollnagel et al., 1999).*

**Fig. S3:**
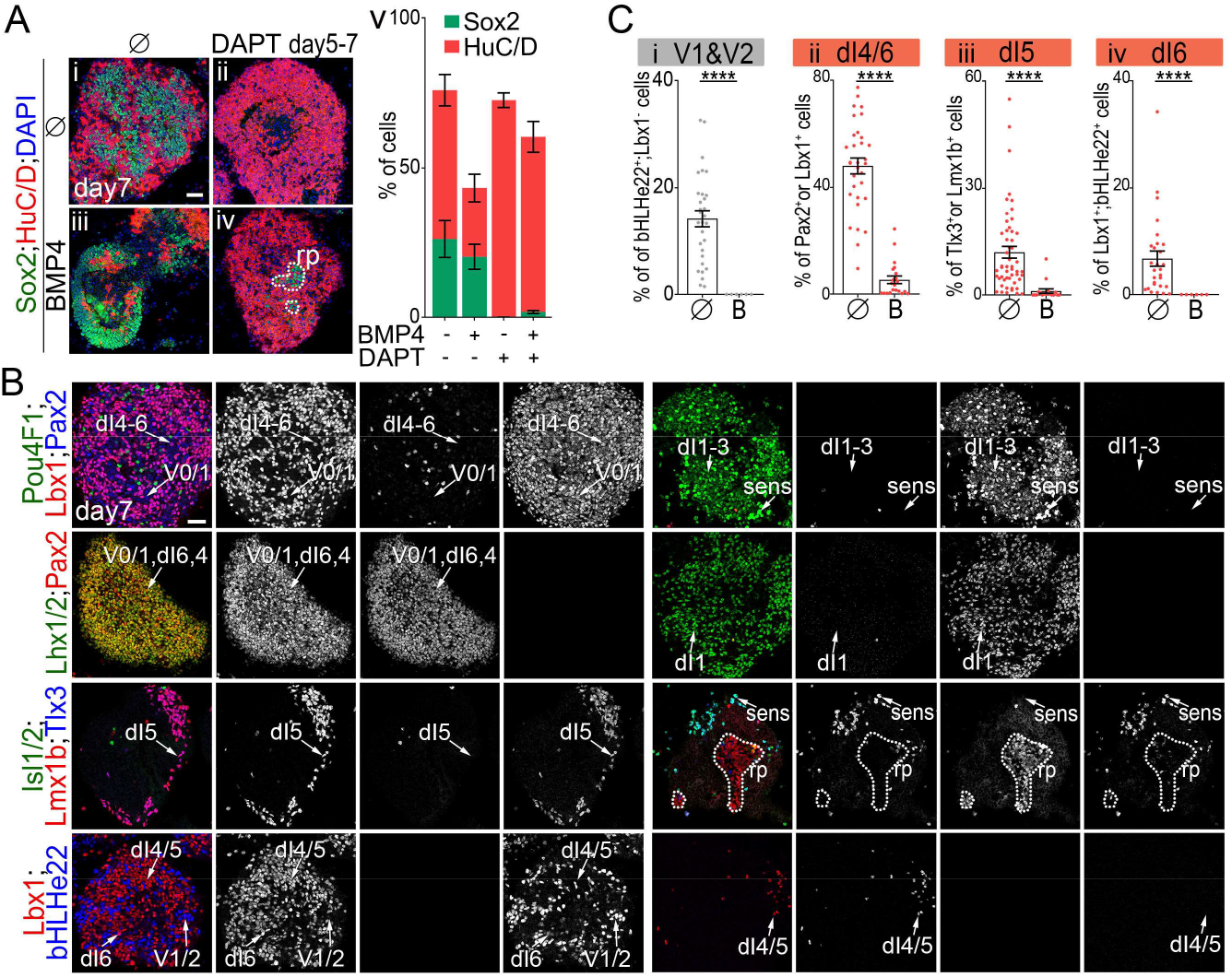
Temporal and spatial dynamics in NP markers expression within dorsal spinal organoids. **(A)** Immunostaining for Sox2 and HuC/D on day 7 EB cultured as in Fig. 1A treated with or without DAPT and with or without BMP4 (i-iv, scale bar: 60μM). Percentage of HuC/D+ (red) and Sox2+ (green) cells in day 7 mouse ESC derived EB grown in the indicated conditions (mean±s.e.m.)(v). *DAPT treatment from day 5 pushed BMP4 treated or not treated cells towards a terminally differentiated HuC/D+ state. BMP4 decreased the rate of differentiation of NP in EB.* **(B)** Immuno-detection for the indicated IN subtype markers on day 7 organoids treated or not with BMP4. Dash lines surround rp territory. Scale bar: 60μM. **(D)** Percentage of ventral and associating IN subtypes harbouring the indicated TF code per image field (circles: individual values, bars: mean±s.e.m.). *In absence of BMP4, the vast majority of cells within the organoid harboured a molecular identity reminiscent of one of the three major classes of early born associating neurons. It included 5% of Lbx1+;bHLHe22+ dI6 cells, 12% of Lbx1+; Pou4f1+ or Tlx3+;Lmx1b+ peripheral dI5 cells, and 45% of Lbx1+;Pax2+;bHLHe22^−^ like cells. The remaining 20% of cells displayed a ventral V2 to V0 IN states. In presence of BMP4 these cell-types are no longer produced in the EB. Instead, a variety of dorsal cells were present in the organoids.*

**Fig. S4:**
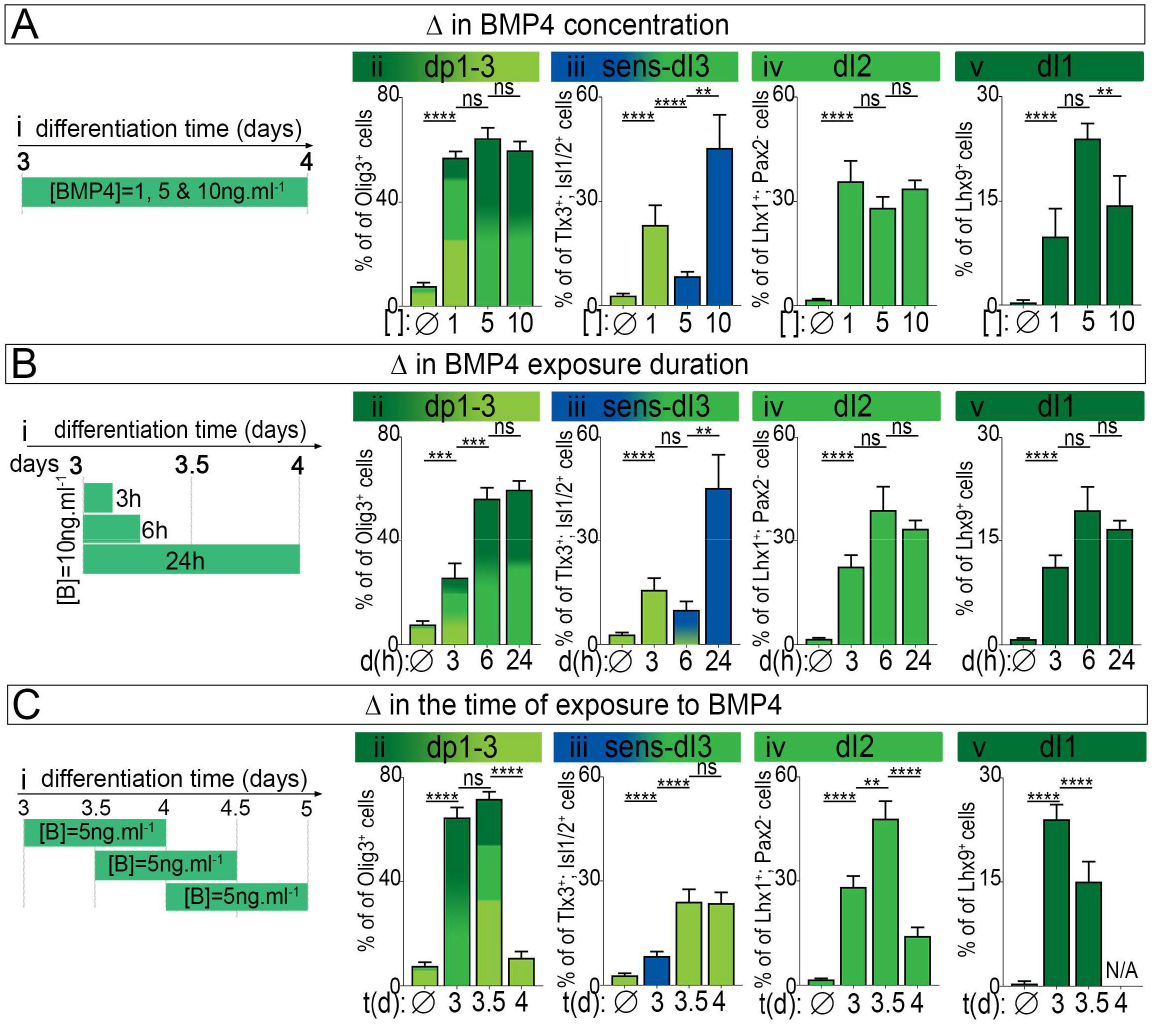
Proportion of relay IN produced upon the modulation of BMP4 concentration, exposure duration and timing. Quantification of the percentage per image field of dp1-dp3 Olig3^+^ NP, dI3, dI2, dI1 and ncc derived sensory cells coloured-coded as in Fig. 1A (circles: individual values, bars: mean±s.e.m.) upon the modulation of BMP4 concentration **(A)**, BMP4 exposure duration **(B)** and the timing of exposure **(C)**. In i, schematics showing the time period and the concentration of BMP4 used. *Increasing BMP4 concentration, duration exposure and advancing the time at which BMP4 is added promotes the generation of cells with a more dorsal identity.*

